# KoRV-related retroviruses in diverse Australian and African rodent species

**DOI:** 10.1101/2024.02.25.581998

**Authors:** Joshua A. Hayward, Gilda Tachedjian

**Author notes:** Corresponding Authors &.

## Abstract

The koala retrovirus (KoRV) is a key contributor to the ongoing decline of Australia’s koala population. KoRV has only been found in koalas and its enigmatic origins, as well as that of its close relative, the gibbon ape leukemia virus (GALV), have been a source of enduring debate. Bats and rodents are each proposed as major reservoirs of interspecies transmission with ongoing efforts to identify additional animal hosts of KoRV-related retroviruses. In this study we identified nine rodent species as novel hosts of KoRV-related retroviruses. Included among these hosts are two African rodents, revealing the first appearance of this clade beyond the Australian and Southeast Asian region. One of these African rodents, *Mastomys natalensis*, carries an endogenous KoRV-related retrovirus that is fully intact and potentially still infectious. Our findings suggest that rodents are the major carriers of KoRV-related retroviruses, with a potential point of origin in Southeast Asia.

## Introduction

Australia’s koala population is in perilous decline (1). Apart from habitat loss, one of the major contributors linked to this decline is disease associated with the koala retrovirus (KoRV), which is widespread in koala populations particularly in Northeastern Australia (2–5). KoRV contributes to koala deaths through its association with diseases including chlamydia, the development of neoplastic lymphoma and immune modulation (6–12).

The origin of KoRV and its close viral relatives have been a topic of sustained interest among the research community (13–24). This is in part because the closest relative of KoRV, the gibbon-ape leukemia virus (GALV) (16, 20), was hosted by captive gibbons whose territory in Southeast Asia does not overlap with koalas in Australia (25). In recent years, bats and rodents have become the prime suspects as transmitters of KoRV-related retroviruses and the origin of KoRV (13, 14) with multiple factors implicating these hosts. First, KoRV-related retroviruses were discovered in taxa other than koalas and gibbons in recent years; initially in an Australian rodent with subspecies in Indonesia (26, 27), and then in Australian and Asian bats whose geographic ranges link those of gibbons and koalas (15, 21). Second, rodents and bats together comprise almost half of all mammalian species, and phylogenomic analyses have revealed that both bats and rodents are significantly involved in the history of retrovirus transmission between other mammalian species (22, 28).

The phylogenetic tree of KoRV and KoRV-related retroviruses remains fragmented, with clear evolutionary gaps (15). In particular, the evolutionary distance between KoRV, GALV, and other viral relatives is still large enough that there are almost certainly other virus-host associations that remain undiscovered. When considering the potential for zoonotic transmission and pathogenicity in humans or other animals of domestic, economic, and ecological importance, questions regarding the origin, transmission, and hosts of KoRV and KoRV-related retroviruses need to be addressed.

Like many other retroviruses, KoRV and GALV are oncogenic, causing blood cancer in koalas and gibbons, respectively (17, 29). Recently, a KoRV-related retrovirus was identified in a fruit bat with lymphoid leukemia (30). Importantly, that virus, the Hervey pteropid gammaretrovirus (HPG), was previously shown to be capable of infecting human cells *in vitro* (15). It remains an open question whether HPG or other KoRV-related retroviruses can establish an infection in humans.

Endogenous ‘fossil’ retrovirus sequences are ubiquitous within the genomes of mammals (31). This is a result of the hallmark of retrovirus replication where the retroviral proviral DNA precursor is inserted into the genome of the host (31). When this happens in germline cells that become new offspring, the retrovirus becomes an endogenised, heritable genetic element. Over the course of evolutionary history, vertebrate genomes have become littered with the remains of past retroviral infections (31). Endogenous retroviruses are subject to genetic drift and tend to become defective over many host generations (31). KoRV subtype A (KoRV-A) is a recently integrated endogenous retrovirus in the koala gene pool, and still generates infectious viral particles that can be transmitted between animals (4, 32). Other variants of KoRV (KoRV subtypes B-M) are understood to circulate among koalas as exogenous retroviruses (3, 33, 34).

Here, we use the term ‘KoRV-related retroviruses’ to describe the endogenous and exogenous retroviruses which form a monophyly with KoRV, and which are not basal to or include the more distantly related *Mus caroli* endogenous retrovirus (McERV) (15). Viruses within this clade may also be referred to as ‘GALV-like’ retroviruses (35). This clade includes, among others, the *Melomys burtoni* retrovirus (MbRV), Melomys woolly monkey retrovirus (MelWMV), as well as the newly reported complete Melomys woolly monkey retrovirus (cMWMV) hosted by Australian and New Guinean *Melomys* rodents, and HPG and the flying fox retrovirus (FFRV1) hosted by Australian bats (15, 26, 27, 36).

We leveraged the vast amount of publicly available data in nucleotide sequence read archives (SRA) and genome assemblies to identify new KoRV-related retroviruses.

## Results

To identify previously unreported KoRV-related retroviruses within publicly accessible datasets, it was necessary to employ a strategy that would distinguish the target retroviral sequences from non-target retroviral sequences that would appear in our search results due to the presence of conserved sequence regions. Additionally, searches with large sequences, such as the complete KoRV genome, are computationally intensive and time consuming to run on public servers. Mammalian SRA records are too large and numerous to make comprehensive local analyses practical for all researchers. To account for these challenges, we used an iterative search strategy where our initial search query employed a short sequence from the least conserved region of the retroviral genome, the receptor binding domain within the *env* gene of a KoRV-related retrovirus. The rationale for this approach is that since there is a large degree of sequence diversity in the receptor binding domain, even among closely related retroviruses (15, 37, 38), any positive hit would likely be to a very close retroviral relative.

### KoRV-related retroviruses were identified in seven Australian rodent species

To identify KoRV-related retroviruses in all Australian mammals for which unassembled SRA have been made public, we searched for available SRA records for 80 species of Australian bats, 69 species of Australian rodents, 165 species of Australian monotremes and marsupials, and 8 species of other Australian eutherian mammals (SI Table 1). For many species, no publicly available SRA records exist. This was acute for Australian bats, with only 11 of 80 species (14%) represented. For other mammal groups, most species were represented among the SRA (SI Table 1): Rodents, 51/66 species (77%); Monotremes and marsupials, 130/165 species (79%); and other eutherian mammals, 8/8 (100%). Using a KoRV-related retroviral receptor binding domain sequence as the search query, positive hits were identified for one bat, one marsupial, and seven rodents (SI Table 1).

Consistent with previous findings, positive hits from the bat, *Pteropus poliocephalus* (Grey-headed flying fox) matched the known KoRV-related retrovirus, HPG (15, 30), and the marsupial, *Phascolarctos cinereus* (koala), was positive for KoRV. Positive hits representing novel retroviral sequences were from the rodents *Mastacomys fuscus* (Broad-toothed rat)*, Pseudomys apodemoides* (Silky mouse)*, P. bolami* (Bolam’s mouse)*, P. delicatulus* (Delicate mouse)*, P. johnsoni* (Central pebble-mound mouse)*, P. shortridgei* (Heath mouse), and *Zyzomys argurus* (Common rock rat). These rodents occupy diverse regions across the Australian continent that collectively include all Australian states and the Northern Territory (Figure 1). SRA reads from each species were extracted and *de novo* assembled into contigs (∼200-1000 nt) for subsequent alignment and phylogenetic analyses (Table 1A). Some contigs from individual species were found to overlap. Overlapping contigs likely represent either different retroviruses or multiple germline insertions/duplications of the same retrovirus whose sequences have diverged over time (31). Contigs whose most similar match was a KoRV-related retrovirus, as determined by BLAST, were selected for subsequent phylogenetic analyses (Figure 2, SI Figures 1-7). Overlapping contigs from individual species were included in the same phylogenies, while non-overlapping contigs were analysed in separate phylogenies. Phylogenies for *gag, pol,* and *en*v retroviral sequences from each rodent species are included in SI Figures 1-4. Additional phylogenies for several of these species are included in SI Figures 5-7. Most contigs assembled from rodent SRA contained deleterious mutations such as frameshifting indels or premature stop codons (Table 1A). This indicates that these contigs were derived from defective endogenous retroviruses. Only one species, *P. delicatulus*, yielded contigs that did not contain deleterious mutations (Table 1A).

**Table 1.**
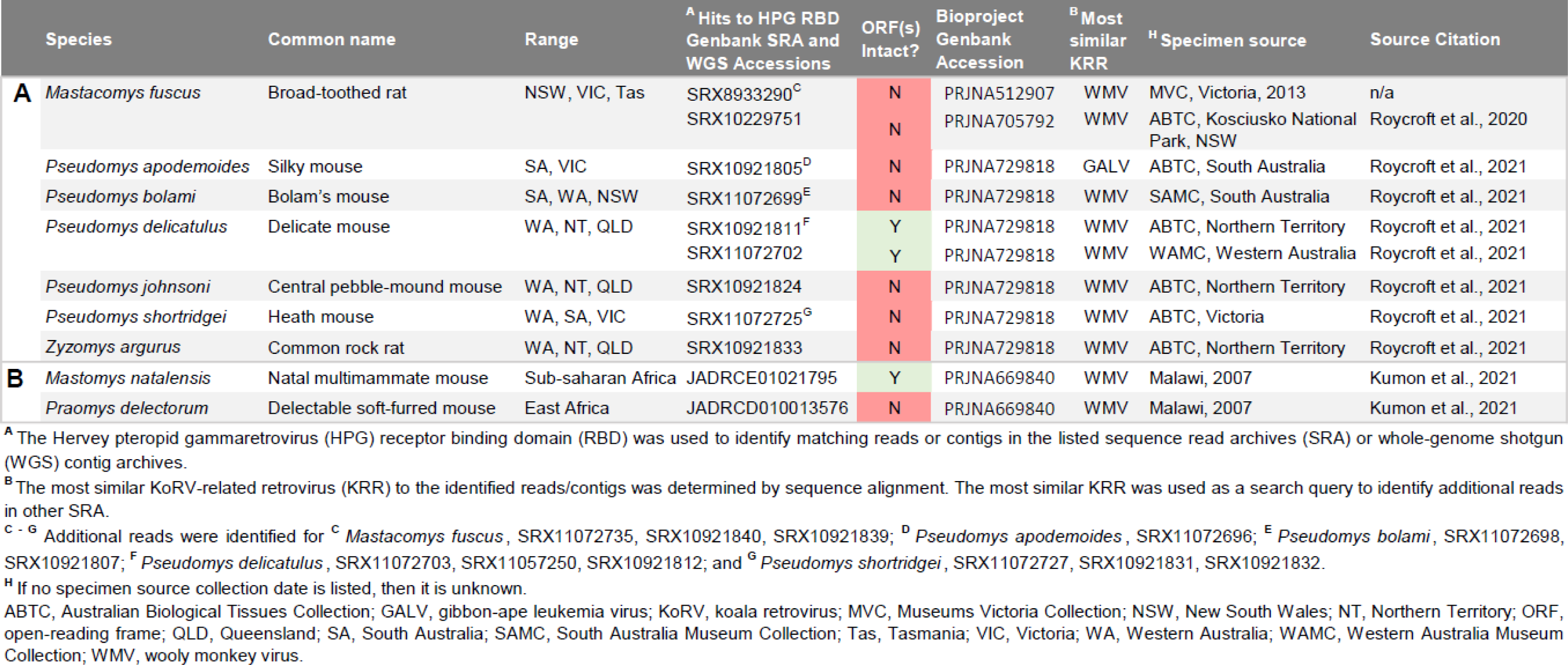
Mammalians species samples in which novel KoRV-related retroviruses were identified using the Hervey pteropid gammaretrovirus receptor binding domain nucleotide sequence.

**Figure 1.**
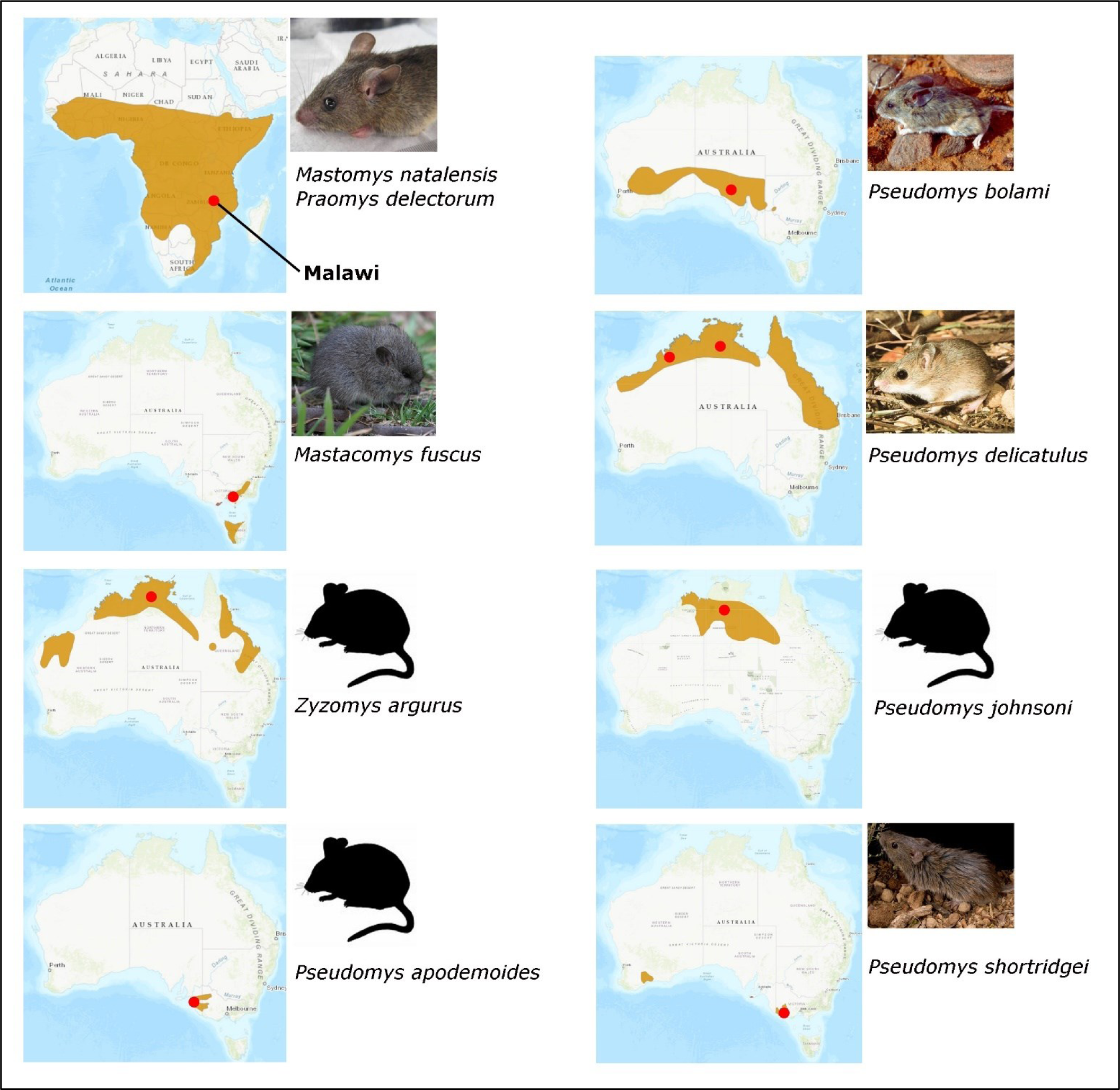
Rodent hosts of KoRV-related retroviruses. Australian and African rodents newly identified in this study as hosts of KoRV-related retroviruses are pictured. The natural ranges of these rodents are indicated by the brown shaded region overlayed on the map of Africa or Australia (59–67). The approximate locations of the collected samples are indicated by the red dots. See the ‘Image Attributions’ section for license details.

**Figure 2.**
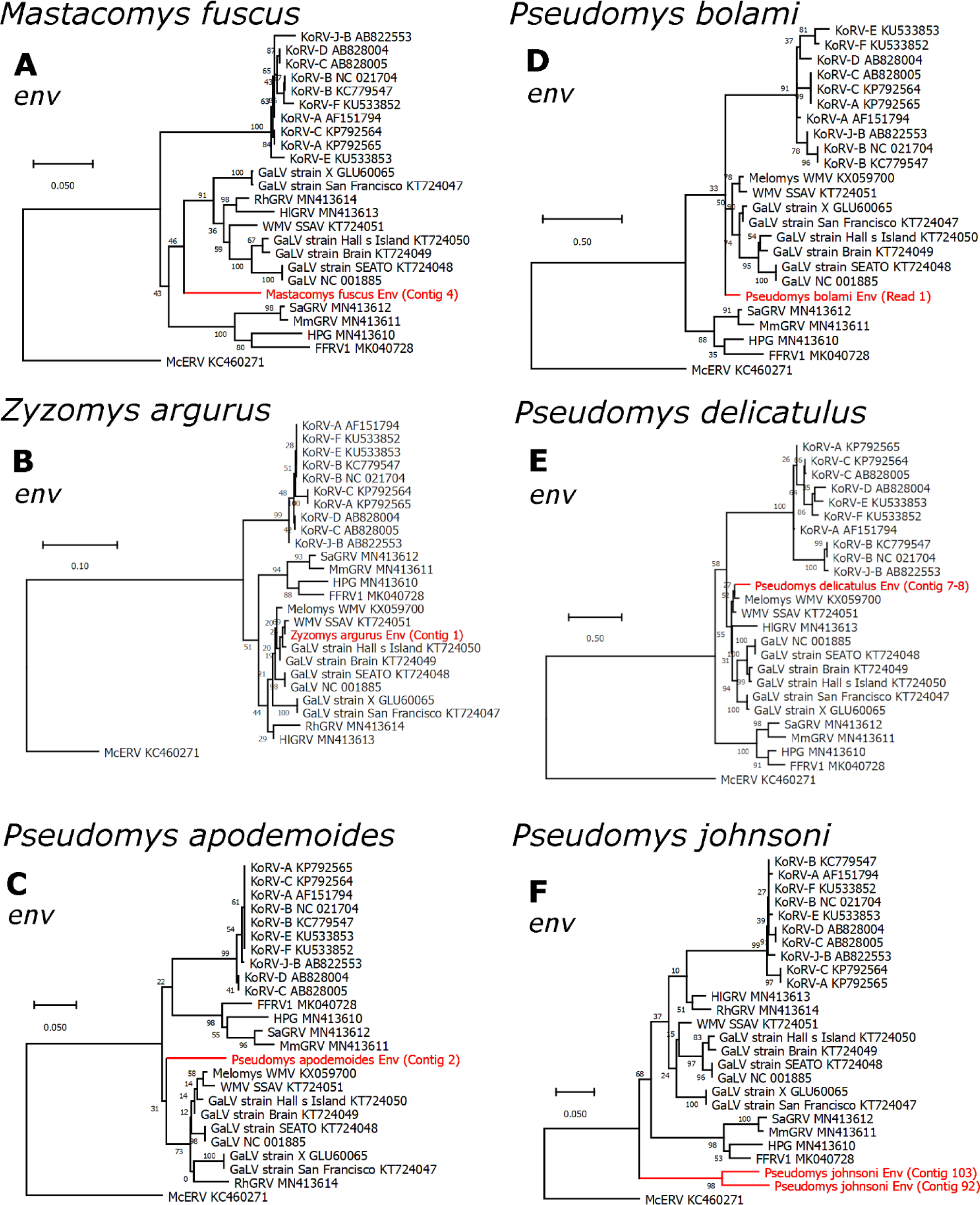
Phylogenetic evolutionary analysis of Australian rodent KoRV-related retroviral sequences. Maximum likelihood phylogenies of regions of nucleotide sequences from the *env* genes of **A**, *Mastacomys fuscus*; **B**, *Zyzomys argurus*; **C**, *Pseudomys apodemoides*; **D**, *Pseudomys bolami*; **E**, *Pseudomys delicatulus*; and **F**, *Pseudomys johnsoni*. All branches are scaled according to the number of nucleotide substitutions per site as indicated by the scale bars. Trees were rooted using the McERV (*Mus caroli* endogenous retrovirus) KC460271. Bootstrap support values are shown at the nodes.

All Australian rodent datasets contain multiple contigs from one or more of the core retroviral genes (*gag*, *pol*, and *env*) that phylogenetically clustered within the KoRV-related retrovirus clade. (Figure 2, SI Figures 1-7). No contigs were identified which were closely and immediately basal to the KoRV sub-clade. For *env* sequences (Figure 2), contigs clustering within the GALV/WMV sub-clade were derived from *Z. argurus* and *P. delicatulus* (Figure 2B & E). *Env* contigs adjacent to the GALV/WMV sub-clade were derived from *M. fuscus*, *P. bolami*, *P. apodemoides* (Figure 2A, C, & D), and *P. shortridgei* (SI Figure 4C).*Env* contigs basal to all other KoRV-related retroviruses were derived from *P. johnsoni* (Figure 2F). For *gag* and *pol*, contigs clustered at variable positions within the phylogeny, including in some cases, at a position basal to McERV (SI Figures 1-7). Together, these contigs may represent multiple divergent gammaretroviruses, or individual recombinant retroviruses that include one or two genes derived from a KoRV-related retrovirus, and the remainder from a more distantly related gammaretrovirus. Taken together these data implicate three novel genera of Australian rodents (*Pseudomys, Mastacomys,* and *Zyzomys*) as hosts of KoRV-related retroviruses, in addition to the *Melomys* genus

### African rodents harbour KoRV-related retroviruses, one of which is intact and potentially infectious

We extended our search for unreported KoRV-related retroviruses to all parts of the world, searching within all mammalian species for which genome assemblies has been made public. We searched the reference sequence (RefSeq) and whole genome shotgun (WGS) assemblies available in GenBank. Searches were performed for high-level taxa representing the breadth of the class Mammalia. Positive hits were identified for numerous taxa within the Laurasiatheria (e.g., bats) and Eurachontoglires (e.g., rodents) superorders, while no hits were identified within primates, Afrotheria, Xenarthra, Marsupialia (except koalas), or Monotremata (SI Table 2). The RefSeq/WGS contig hits were extracted and annotated for subsequent analyses (Table 2).

**Table 2.**
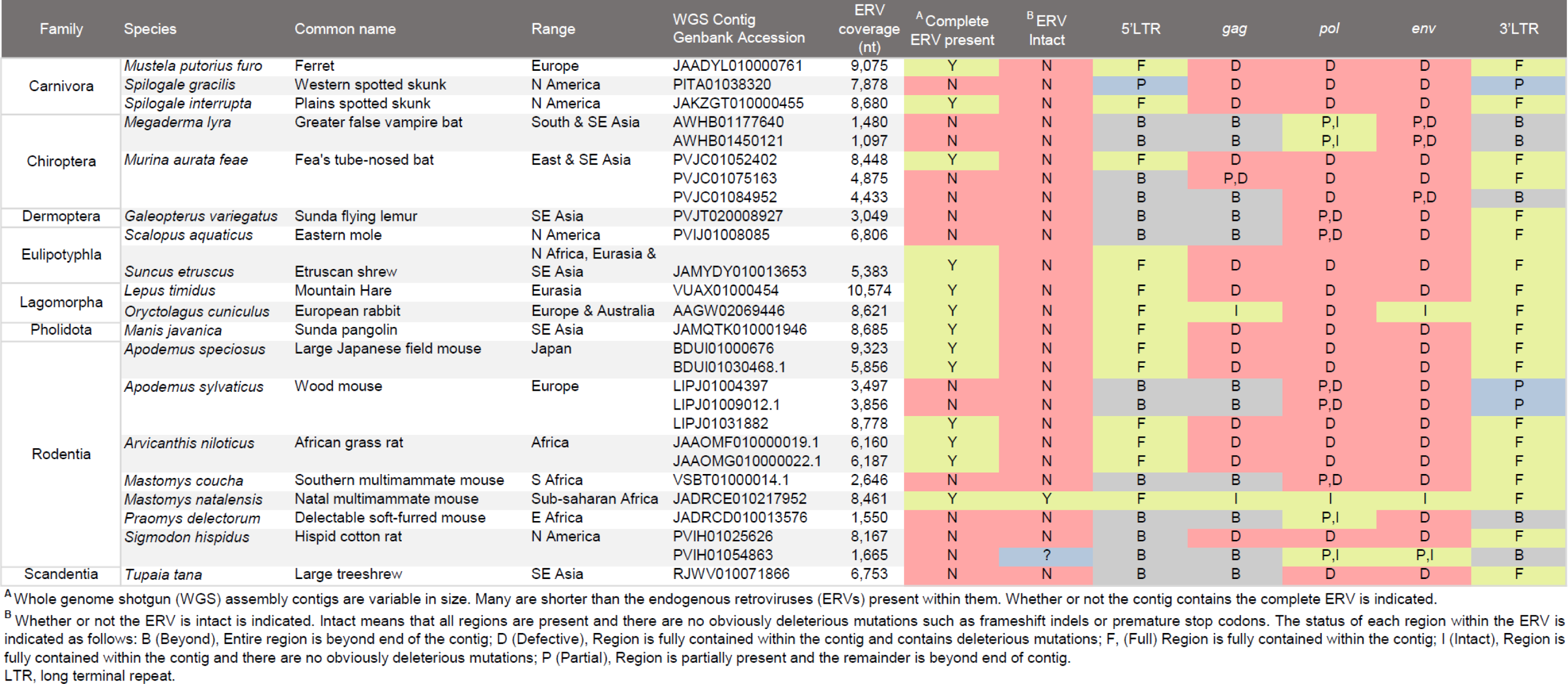
Mammalian ERVs identified in WGS assemblies using the Hervey pteropid gammaretrovirus receptor binding domain nucleotide sequence.

The sequences identified in genome assemblies represent more complete (1,097 – 10,574 nt) endogenous retroviruses as they were extracted from larger genome assembly contigs compared to the short contigs assembled for the Australian rodents from SRA. Among the endogenous retroviruses identified in this search, two were found to phylogenetically cluster within the KoRV-related retrovirus clade, (Figure 3, Tables 1B and 2). These KoRV-related retroviruses are hosted by the African rodents, *Mastomys natalensis* and *Praomys delectorum*.

**Figure 3.**
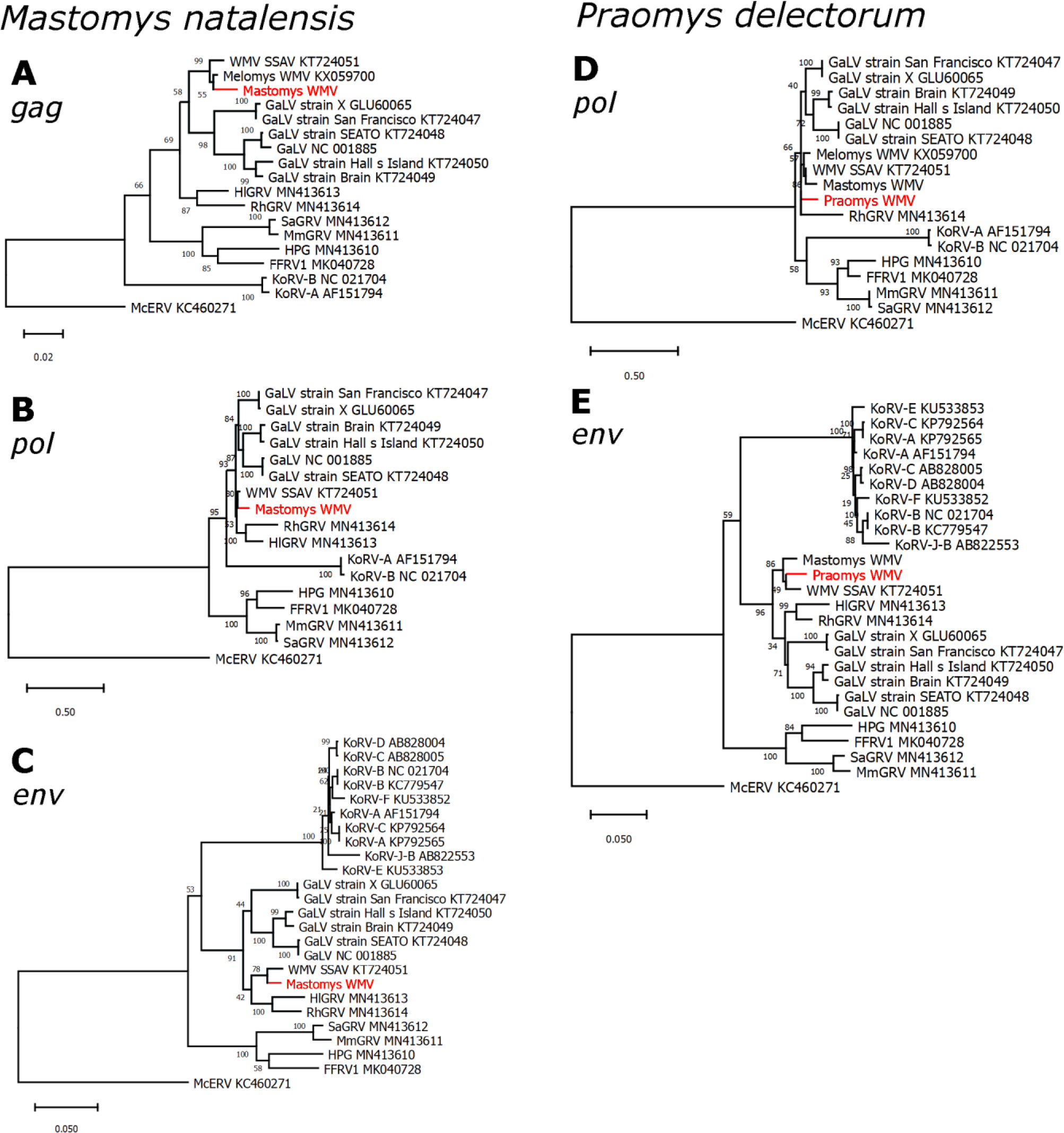
Phylogenetic evolutionary analysis of African *Mastomys natalensis and Praomys delectorum* KoRV-related retroviruses. Maximum likelihood phylogenies of the A, *gag*, B, D *pol*, and C, E *env* genes of KoRV-related retroviruses and the novel **A-C** Mastomys WMV (*Mastomys* woolly monkey virus) and **D-E** Praomys WMV. All branches are scaled according to the number of nucleotide substitutions per site as indicated by the scale bars. Trees were rooted using the McERV (*Mus caroli* endogenous retrovirus) KC460271 sequences. Bootstrap support values are shown at the nodes.

The first African KoRV-related retrovirus, derived from *Mastomys natalensis* and designated as Mastomys WMV, is completely intact (Figure 4) and has a high 96.9% nucleotide sequence identity to the woolly monkey simian sarcoma virus (WMV SSAV), a KoRV-related retrovirus. For comparison, Mastomys WMV shares 79.9% and 88.7% nucleotide identity with KoRV-A and GALV, respectively. Mastomys WMV contains the expected open reading frames for *gag, pol* and *env* and all the expected functional motifs are conserved (Figure 4). These include: the proline primer binding site and polyadenylation signal, the major homology region and CCHC zinc finger of Gag, and the protease, reverse transcriptase, and integrase enzymatic active sites (Figure 4). In addition, many of the nucleotide differences in the receptor binding domain compared to WMV SSAV are non-synonymous mutations, and the CETTG pathogenicity motif is present in *env* (Figure 4 & SI Figure 8).

**Figure 4.**
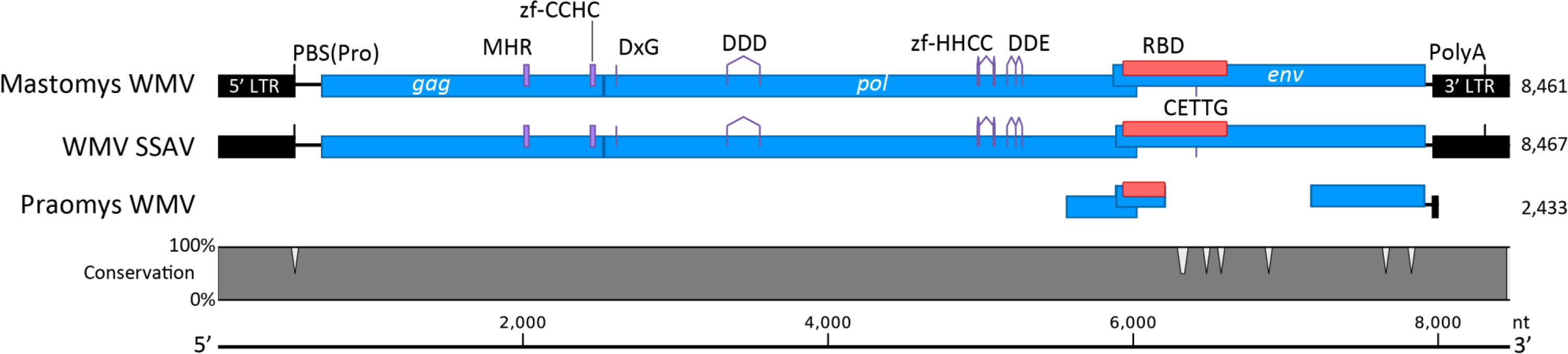
Alignment of the Mastomys WMV, Praomys WMV, & WMV SSAV. Open reading frames encoding the core retroviral genes *gag, pol,* and *env*, (blue regions) and the 5’ and 3’ long terminal repeats (LTR; black regions) are depicted. The alignment scale is in nucleotides and the total length of each sequence is listed on the right side of the alignment. The Conservation line graph (grey region) depicts nucleotide mismatches between Mastomys WMV and WMV SSAV. Conserved functional motifs (purple regions) are indicated: PBS(Pro), proline tRNA primer-binding site; MHR, major homology region; zf, zinc finger; DxG, protease active site motif; DDD, reverse transcriptase active site motif; DDE, integrase active site motif; RBD (red regions), receptor binding domain; CETTG, pathogenicity motif; PolyA, polyadenylation signal; *env*, envelope; *gag*, group-specific antigen; Mastomys WMV, *Mastomys* woolly monkey virus; WMV SSAV, woolly monkey simian sarcoma virus; *pol*, polymerase.

The 5’ and 3’ long terminal repeats (LTRs) of Mastomys WMV are 100% identical (Figure 4). The lack of nucleotide differences resulting from genetic drift since integration indicates that this retrovirus integrated into the genome of *M. natalensis* recently. The presence of intact *gag*, *pol*, and *env* genes, and functional enzymatic motifs suggests that Mastomys WMV may be capable of expressing infectious viral particles.

The second African KoRV-related retrovirus, derived from *Praomys delectorum*, and designated as Praomys WMV, was identified on a relatively short genomic contig containing only the 3’ end of the *pol* gene, and the 5’ and 3’ ends of the *env* gene with an 889 nt internal deletion relative to WMV SSAV (Figure 4). The remainder of the *env* gene, and the partial *pol* gene present on this contig contain mostly uninterrupted reading frames. This indicates that this is a defective endogenous retrovirus. It shares 95.2% nucleotide identity with WMV SSAV. Both Mastomys WMV and Praomys WMV phylogenetically cluster very closely with WMV SSAV and another closely related, defective Australian rodent retrovirus, Melomys WMV (Figure 3).

### Novel hosts of KoRV-related retroviruses possess the PiT-1 cell receptor permissivity motif

A commonality among most KoRV-related retroviruses is their apparent shared use of the PiT-1 (SLC20A1) cellular receptor (15, 37, 39, 40). The use of PiT-1 for viral entry requires the presence of a permissive protein sequence motif (15, 41, 42). In contrast *Mus musculus* does not possess this motif and is not susceptible to KoRV-related retroviruses (15). We analysed the PiT-1 sequences for all the novel hosts of KoRV-related retroviruses reported here. All but one rodent PiT-1 sequence from *Zyzomys argurus*, which contains a codon deletion, appear to possess the permissive motif (SI Figure 9). This suggests that these rodents are likely susceptible to cell entry by KoRV-related retroviruses engaging the PiT-1 receptor, with the possible exception of *Zyzomys argurus* (SI Figure 9).

### Phylogenetic analysis of clade-adjacent endogenous retroviruses suggests a potential Southeast Asian rodent origin for KoRV-related retroviruses

Phylogenetic analysis of the endogenous retroviruses extracted from RefSeq/WGS genome assemblies revealed that most of the endogenous retroviruses identified through sequence homology with the HPG receptor binding domain were gammaretroviruses outside the KoRV-related retrovirus sub-clade (Figure 5). This analysis also revealed a geographic bias among the hosts of these endogenous retroviruses (Figure 5 & Table 2). Using our targeted search strategy, 27 endogenous retroviruses were identified across 19 species (Table 2). Of these, six species are present in Southeast Asia, and one in Australia, comprising 37% of the host species. These Southeast Asian host species are diverse and include microbats (*Megaderma lyra* and *Murina aurata feae*), lemurs (*Galeopterus variegatus*), Etruscan shrews (*Suncus etruscus*), pangolins (*Manis javanica*), and treeshrews (*Tupaia tana*). Further, among the identified host taxa, 7 of the 19 (37%) species are rodents (Table 2). These data indicate that despite not phylogenetically clustering within the KoRV-related retrovirus sub-clade, a large proportion of gammaretroviruses with receptor binding domain sequence homology to KoRV-related retroviruses are hosted by Southeast Asian mammals and the host species are predominantly rodents.

**Figure 5.**
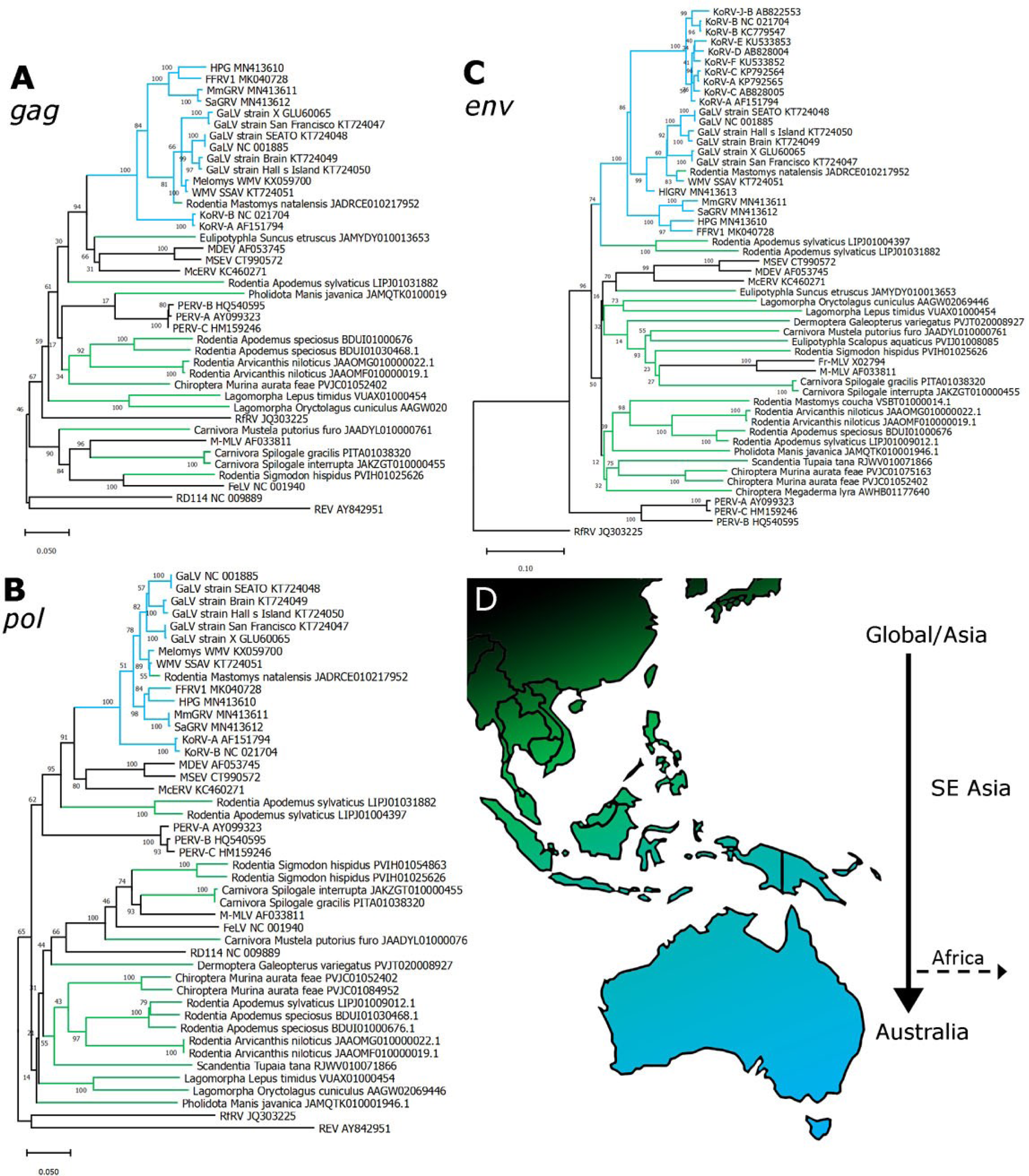
Evolutionary relationships of novel endogenous gammaretroviruses identified in mammalian genome assemblies via sequence homology with a KoRV-related retroviral receptor binding domain sequence. Maximum likelihood phylogenies of the A *gag*, B *pol*, and C *env* genes. All branches are scaled according to the number of nucleotide substitutions per site as indicated by the scale bars. Branches representing the KoRV-related retrovirus clade are shown in blue; branches representing novel endogenous retroviruses reported in this study are shown in green; branches representing previously reported gammaretroviruses are shown in black. The dashed arrow represents the hypothetical recent transmission from Australia/SE Asia to Africa suggested by the presence of KoRV-related retroviruses in the genomes of African rodents. Bootstrap support values are shown at the nodes. **D** Schematic representation of the Southeast Asian/Australian region. The arrow, and colour gradient (Black, Global/Asia; Green, SE Asia; Blue Australo-Papuan region) indicates the hypothetical average direction of transmission of KoRV-related retroviruses through evolutionary history.

## Discussion

KoRV and its close relatives are viruses of ecological concern, with origins shrouded in mystery, and potentially pathogenic consequences in the event of further cross-species transmission events into humans or other animals. To enhance our understanding of the breadth of the host network of this group of viruses and identify hosts which may require further attention, we searched for previously unreported KoRV-related retrovirus sequences hidden in the growing expanse of publicly available sequence data.

### KoRV-related retroviruses in multiple species of Australian rodents

We identified KoRV-related retroviral sequences in seven species of Australian rodents (Table 1A). All SRA records in which these retroviral sequences were identified were exomes recently generated from Australian museum specimens (43, 44). These rodent species inhabit diverse territories around much of coastal Australia, including Tasmania (Figure 1). KoRV itself is present in koalas from Queensland, New South Wales, and to a lesser extent in Victoria and South Australia (3). However, reported rodent KoRV-related retroviruses in Australia had previously been limited to the host *Melomys burtoni,* which ranges in the north and northeastern coast of Australia (45). This geographic distribution suggests that KoRV and KoRV-related retroviruses are widespread across Australia and may infect many more hosts than previously known.

Given the limited read depth of the rodent SRA datasets only short contigs could be assembled from reads with sequence homology to KoRV-related retroviruses. Despite this limitation, deleterious mutations in these retroviruses were sufficiently abundant that almost all contigs were found to contain them. This suggests that they are likely endogenous retroviruses and that their integration was not recent. Complete endogenous retroviral genomes from these rodents will help to determine how long ago these integration events occurred.

Many of the assembled contigs overlapped in their positions of alignment to KoRV-related retroviral genomes. In these cases, they could be included in the same phylogenies (Figure 2F, SI Figures 1-3, 5-7). Overlapping but distinct contigs indicate the presence of either multiple retroviruses, or duplication of a single integrated retrovirus followed by sequence divergence (31). Clarifying this will require more extensive sequencing of these rodent genomes. It is also worth noting that five of the seven Australian rodent species are members of the same genus, *Pseudomys*. Together with the observation that these retroviral sequences? are likely not recent integrations, this might suggest that one or more of the endogenous retroviruses represented by these contigs integrated into a common ancestor of these rodents.

For each Australian rodent species, the KoRV-related retroviral contigs representing the *gag, pol*, and *env* genes, appeared at varying positions in the phylogenies relative to the KoRV, GALV, WMV, and bat virus sub-clades (Figure 2, SI Figures 1-7). This may indicate that the analysed *gag, pol*, and *env* contigs are derived from different endogenous retroviruses, or that the limited sequence information in these relatively short contigs prevents robust estimation of evolutionary relationships. The latter possibility is suggested by the weak bootstrap support of some internal nodes across the phylogenies (Figure 2, SI Figures 1-7).

### Identification of KoRV-related retroviruses in African rodents

Surprisingly, we identified KoRV-related retroviruses in the genomes of two African rodents. To our knowledge this is the first report of KoRV-related retroviruses infecting hosts outside of Australia and Southeast Asia. Both endogenous retroviruses appear to be very recently integrated into the *Mastomys natalensis* and *Praomys delectorum* genomes. This is evidenced by the 100% nucleotide identity between the Mastomys WMV long-terminal repeat regions, and a very low frequency of indel mutations in Praomys WMV.

Mastomys WMV has a fully intact proviral genome. Previously reported KoRV-related retroviruses from rodents have contained incomplete or otherwise defective genomes. The only exception to this is cMWMV, which is a newly reported, infectious, non-fixed endogenous KoRV-related retrovirus present in a subset of individuals of the New Guinea rodent species *Melomys leucogaster* (35). All canonical genes are present in Mastomys WMV, with uninterrupted open reading frames and the conservation of expected functional motifs (Figure 4). The presence of multiple non-synonymous mutations in the receptor binding domain is consistent with ongoing selective pressures in different hosts following the divergence of WMV SSAV and Mastomys WMV from their common ancestor. If rodents are the major hosts of this clade of retroviruses and responsible for many interspecies transmission events, then the identification of potentially infectious rodent KoRV-related retroviruses fills an important gap in supporting this notion.

Mastomys WMV may reflect a similar case to KoRV-A and cMWMV in which the virus, being recently integrated, is in the process of endogenisation and fixation in the gene pool while still producing functional, infectious retroviral particles. Extended sampling and analysis of Mastomys WMV presence among individual *Mastomys natalensis* rodents may clarify this possibility. Praomys WMV may also be in the process of undergoing endogenisation and fixation in its murine host; however, the identified copy is clearly no longer functional since it has a single deletion of 889 nt (relative to WMV SSAV) in the *env* gene that would prohibit this endogenous retrovirus from generating infectious viral particles. Mastomys WMV and Praomys WMV appear to be distinct retroviruses, as each has higher nucleotide identity to WMV SSAV than to each other (94.3%), suggesting that they are not the result of a single integration event prior to speciation of the *Mastomys* and *Praomys* hosts.

The high degree of nucleotide identity of these African retroviruses to WMV SSAV and MelWMV suggests recent transmission. Rodents of various species have been documented as pests aboard ships (46, 47). There are shipping routes across the Indian ocean connecting Southeast Asia, Australia, and East Africa, and it seems reasonable to speculate that a rodent stowaway carrying an infectious WMV variant might explain these findings. The newly reported cMWMV shares exceptionally high nucleotide identity with WMV (98.9%) (35), which is closer than the nucleotide identity shared between Mastomys WMV, Praomys WMV, and WMV. Taken together, these studies suggest that the WMV sub-clade is actively undergoing transmission between rodent species/genera and geographic regions.

### Possible Southeast Asian Origins of KoRV-related retroviruses

We searched publicly available genome assemblies for evidence of endogenous retroviruses belonging to the KoRV-related retrovirus clade using a stringent search query comprised of the receptor binding domain of a KoRV-related retrovirus. This did not fully exclude retroviruses from outside this clade from appearing among the positive search hits. Intriguingly, a substantial proportion (37%) of the hosts of the identified retroviruses are found across Southeast Asia and Australia (Table 2). Similarly, many (37%) of these hosts are rodents (Table 2).

Phylogenetic analysis of the non-KoRV-related retroviruses identified by this search strategy provides some insight into their pedigree (Figure 5). While the *gag* and *pol* genes of these endogenous retroviruses are widely dispersed throughout the gammaretroviral phylogeny (Figure 5A & B), the *env* genes cluster within a sister clade to the KoRV-related retroviruses (Figure 5C). This suggests a history of recombination between KoRV-related retroviruses and more distantly related gammaretroviruses. Furthermore, all extant retroviruses within the *env* sister clade are rodent gammaretroviruses such as the Moloney murine leukemia virus (MMLV) and *Mus caroli* endogenous retrovirus (McERV). These are hosted by Asian mice, some of which are now globally distributed.

These data suggest an Asian/Southeast Asian origin for the common ancestor of all KoRV-related retroviruses, with a history of circulation and recombination and transmission into incidental Southeast Asian mammalian hosts, and occasional transmission outside the region. Currently, members of this clade are undergoing active endogenisation in rodent and marsupial hosts in the Australia and New Guinea (35). The presence of KoRV in koalas and WMV variants in numerous Australian and African rodents, suggest one or more transmission events into Australia and, perhaps very recently, into East Africa.

An Important caveat here is the possibility of sampling bias among the genomes represented in the public datasets we queried influencing the apparent abundance of KoRV-related retroviruses in Southeast Asian species relative to other locations. Mammalian species from some geographical regions are likely to be better represented among genome datasets than others. It is worth noting that at present, the RefSeq database (Release 212) contains 1,443 mammalian genomes, which represents only ∼22% of the currently recognised 6,495 mammalian species (48, 49). Future analyses that include a larger, more comprehensive, and geographically unbiased collection of reference mammalian genomes may clarify this possibility. Intriguingly, the new report by Mottaghinia *et al.* (35) suggests an Australo-Papuan origin for this clade (therein referred to as ‘GALV-like’ retroviruses) following the identification of vertically transmitted ERVs in *Melomys* rodents which, as with KoRV, is at the earliest stages of endogenisation. Here we report an additional recent endogenisation in the African rodent *M. natalensis*. The picture of modern KoRV-related retroviral infection and endogenisation is clearly far from complete. Further taxa screening is almost certain to reveal further details of this unfolding story.

These findings support the hypothesis that rodents are the primary host of KoRV-related retroviruses. While other mammals are clearly susceptible to infection with these gammaretroviruses, such transmission events may be incidental to the main thread of gammaretroviral divergence and evolution in rodent hosts. Bats are a potential exception. HPG appears to be endemic among Australian black flying foxes, and HPG’s closest relatives have been found within several different species of pteropid bats. (15, 30). The KoRV-related retroviruses of pteropid bats form a monophyly that does not contain any known rodent viruses (15). This indicates that the HPG sub-clade has become well adapted to circulation among, and transmission between, different bat species. This successful adaptation has not yet been observed for KoRV-related retroviruses infecting other non-rodent clades of mammals.

### Limitations of the study

An important limitation of this study is the lack of breadth of available sequence data. This is somewhat surprising considering that such large amounts of data already exist, but it is important to consider that for most species, data exists from a very limited number of individuals, and for many species no data exists at all. We might compare it to the analogy of trying to find a very specific type of human virus by analysing genomic and transcriptomic data from just a few people. The chances of success might be slim. It’s important to consider this when discussing the host range of KoRV-related retroviruses. While over recent years we have been discovering new host species (15, 21, 26, 27), and this study adds to that number, we may still have only scratched the surface. There may yet be any number of important animal species under threat from this group of cancer-causing viral pathogens.

### Conclusion

The identification of numerous rodents as novel hosts of KoRV-related retroviruses widens our understanding of the host range of this clade of oncogenic viruses. It also highlights that our knowledge remains relatively limited. Many further animal species may already host these viruses or be susceptible to infection. The finding of KoRV-related retroviruses in African rodents, and the expanding association with rodent hosts more generally, demonstrates the transmissibility of these viruses and implicates rodents as an important host reservoir. Since KoRV-related retroviruses are associated with animal disease, future studies should seek to determine which animals could potentially become infected with them. Transmission of KoRV-related retroviruses from rodents or bats into animals of domestic, economic, or ecological importance could have dire consequences, as it has had for koalas.

## Methods

Our use of the term ‘KoRV-related retroviruses’ applies to retroviruses for which at least one of the three gammaretroviral genes *gag*, *pol*, or *env* are within this monophyly. This is because gammaretroviruses are known to undergo recombination (50–53) and consequently may be directly derived from a KoRV-related retrovirus even if one or two of their genes are derived from other more distantly related retroviruses.

### Database searches

To identify KoRV-related retroviruses, we interrogated the available SRA data for each species of interest. The search query was a 540 nt sequence from the receptor binding domain of HPG (SI Data 1). HPG is a KoRV-related retrovirus phylogenetically basal to KoRV and GALV (15) and could be reasonably expected to possess homology to novel KoRV-related retroviruses also basal to the existing monophyly in addition to those phylogenetically intermediate to known KoRV-related retroviruses.

Searches were performed using the Sequence Read Archive Nucleotide BLAST (SRA BLAST) program through the NCBI web interface (https://blast.ncbi.nlm.nih.gov/Blast.cgi?PROGRAM=blastn&BLAST_PROGRAMS=megaBlast&PAGE_TYPE=BlastSearch&BLAST_SPEC=SRA&SHOW_DEFAULTS=on). Search parameters were left in their default settings except for the following: Optimized for ‘Somewhat similar sequences’; Word size = 7; Max target sequences = 1000; Max matches in a query range = 21; and Expect threshold = 1×10^-5^.

To identify KoRV-related endogenous retroviruses within genome assemblies, we used the BLASTn program through the NCBI web interface (https://blast.ncbi.nlm.nih.gov/Blast.cgi) using the same search query as that for the SRA. Search parameters were left in their default settings except for the following: Database was set to ‘refseq_representative_genomes’, ‘refseq_genomes’, or ‘wgs’; Limited by organisms set to each of the ‘Search taxa’ listed in SI Table 2, Optimized for ‘Somewhat similar sequences’; Word size = 7; Max target sequences = 1000; and Expect threshold = 1×10^-20^.

### KoRV-related retrovirus contig assembly

To determine if the closest retroviral match for the identified SRA reads was a KoRV-related retrovirus, a local reciprocal BLAST analysis (54) against known retroviruses was performed. SRA containing reads for which the closest match was a KoRV-related retrovirus were further analysed as follows: SRA BLAST searches used the genome region spanning the *gag*, *pol*, and *env* genes of the closest matching retrovirus as the search query. Search parameters were left in their default settings except for the following: Optimized for ‘Somewhat similar sequences’; Word size = 7; Max target sequences = 1000; Max matches in a query range = 21; and Expect threshold = 1×10^-5^.

Matching SRA reads were collected and underwent *de novo* assembly into contigs using the Assemble Sequences tool in CLC Genomics 22 (CLC; QIAGEN). A local BLAST analysis against known retroviruses was performed for the assembled contigs to identify those whose closest match was a KoRV-related retrovirus.

### Annotation of Mastomys WMV genome

The Mastomys WMV genome sequence was annotated using CLC following alignment using MUSCLE (55), and comparison against the genomes of WMV SSAV (Genbank: KT724051) and HPG (Genbank: MN413610).

### Phylogenetic analyses

To estimate the evolutionary relationships between the endogenous retroviral sequences identified in genome assemblies and SRA with known gammaretrovirus, we conducted phylogenetic analyses. Endogenous retroviruses were extracted from genome assembly contigs by delineation of their long-terminal repeats as described previously (50). The *gag*, *pol*, and *env* genes were identified and annotated by sequence alignment against known gammaretrovirus genes using MUSCLE followed by manual curation. Contigs/reads from SRA were aligned to the *gag*, *pol*, and *env* genes of known gammaretroviruses using the progressive alignment tool in CLC with the following parameters: Gap open cost = 5; Gap extension cost = 2; End gap cost = free; Alignment speed = ‘Very accurate’. Contigs derived from *Pseudomys delicatulus* (*env*), and *P. shortridgei* (*env*), did not overlap any other contigs or reads; in these cases, adjacent contigs were concatenated to improve phylogenetic resolution. Ambiguous regions of the alignments were removed using Gblocks (56).

The best-fit models of nucleotide substitution were determined using the Model Testing tool in CLC, which were found to be GTR+Γ+T for all alignments. Maximum likelihood trees were then inferred using the best-fit model in CLC with 1,000 bootstrap replicates. Trees were visualized with MEGA 11.0 (57).

### PiT-1 sequence analysis

PiT-1 (SLC20A1) genes for species with assembled genomes were extracted from GenBank (SI Table 2). For each of the rodents without assembled genomes, the PiT-1 gene was assembled from the same SRA data set that the KoRV-related retroviral sequences were extracted as follows: The PiT-1 gene of the rodent, *Mastomys coucha* (GenBank: XM_031371865), was used as a search query. An SRA BLAST was conducted with search parameters left in their default settings except for the following: Optimized for ‘Highly similar sequences’; Word size = 16; Max target sequences = 1000; and Expect threshold = 1×10^-20^. Reads were *de novo* assembled in CLC and the resultant contig confirmed to encode PiT-1 by reciprocal BLAST analysis and sequence alignment using MUSCLE.

## Image attributions

Figure 1: “*Mastomys natalensis*, aka the natal multimammate mouse” by JD Kelly et al. (58), available at https://upload.wikimedia.org/wikipedia/commons/1/18/Mastomys_natalensis.jpg under a Creative Commons Attribution 3.0; “Broad-toothed mouse (*Mastacomys fuscus*); Thredbo, NSW, Australia” by M Kjaergaard, available under a Creative Commons Attribution 3.0 at https://upload.wikimedia.org/wikipedia/commons/2/24/Broad_toothed_mouse_1.JPG; “Bolam’s mouse (*Pseudomys bolami*)” by A Stanley, available under a Creative Commons Attribution 4.0 at https://upload.wikimedia.org/wikipedia/commons/4/46/Bolam%27s_mouse_%28Pseudomys_bolami%29_.jpeg; “Little Native Mouse”, by ZooPro, available under a Creative Commons Attribution 3.0 at https://upload.wikimedia.org/wikipedia/commons/0/04/Pseudomys_delicatus_20020401.jpg; “Heath mouse, native Australian rodent at Black Range State Park” by David Paul, available at https://upload.wikimedia.org/wikipedia/commons/8/81/Pseudomys_shortridgei%2C_Heath_mouse_01.jpg under a Creative Commons Attribution 4.0.

## Data availability

GenBank accessions for the publicly available sequence data used in this study are listed in Tables 1 and 2, and SI Table 3.

## Acknowledgements

We thank Edward C. Holmes and Mark Ziemann for their kind assistance and input in drafting the manuscript. J.A.H and this study were funded through the kind support of the Upotipotpon Foundation, the Estate of GWA Griffiths, Miss Gwen Hotton, Mrs Rosemary Jacoby, and the Jim and Margaret Beever Fellowship. We gratefully acknowledge the contribution of the Victorian Operational Infrastructure Support Program, received by the Burnet Institute.

