## Supplementary material for "KoRV-related retroviruses in diverse Australian and African rodent species": SI Figures 1-9

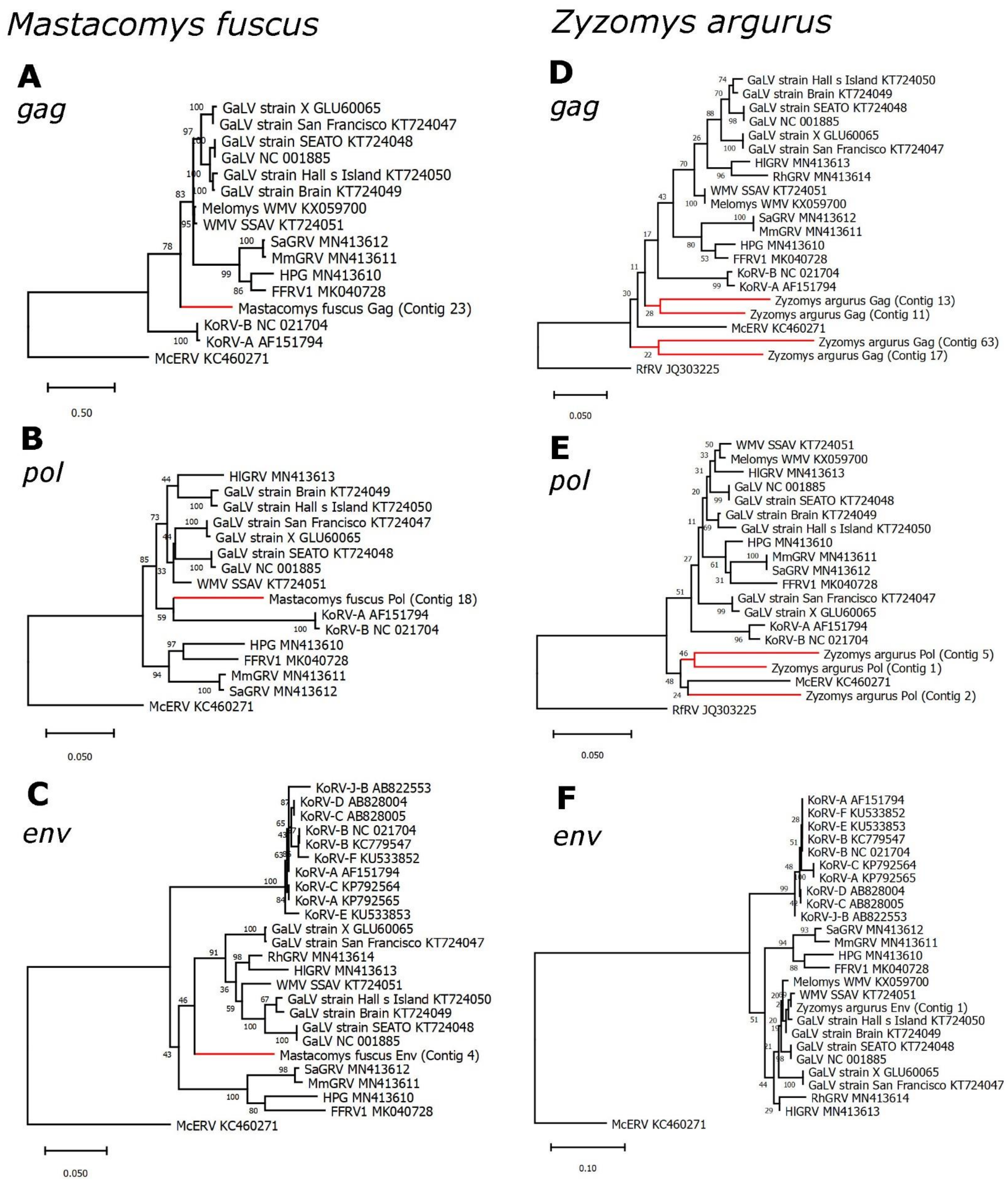


**SI Figure 1. Phylogenetic evolutionary analysis of Australian *Mastacomys fuscus* and *Zyzomys argurus* KoRV-related *gag pol* and *env* retroviral sequences.** Maximum likelihood phylogenies of regions of nucleotide sequences from the **A, D** *gag*, **B, E** *pol*, and **C, F** *env* genes overlapping contigs from **A-C** *M. fuscus* and **D-F** *Z. argurus*. All branches are scaled according to the number of nucleotide substitutions per site as indicated by the scale bars. Trees were rooted using the McERV (*Mus caroli* endogenous retrovirus) KC460271 or RfRV (*Rhinolophus ferrumequinum* retrovirus) JQ303225 sequences. Bootstrap support values are shown at the nodes.

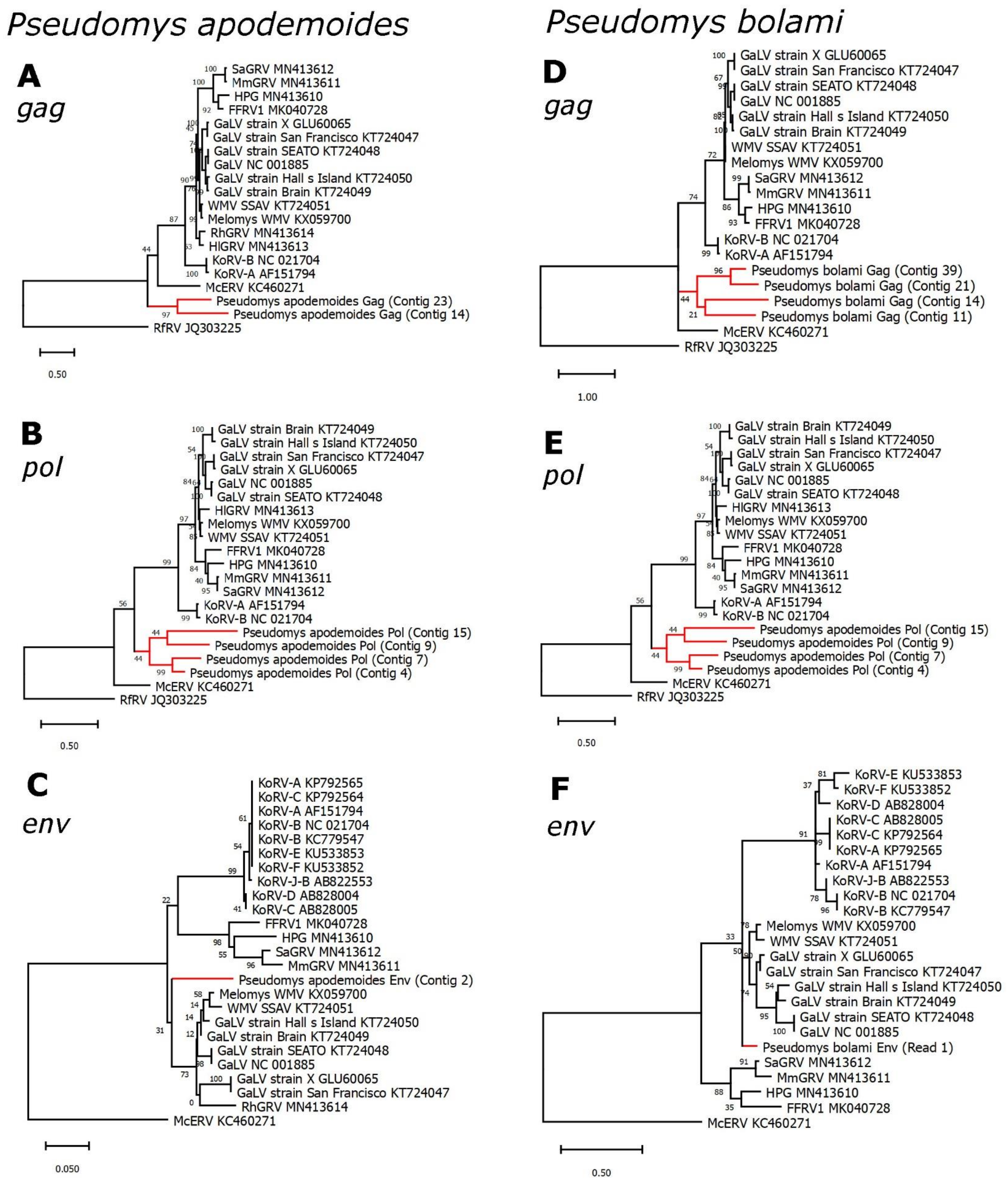


**SI Figure 2. Phylogenetic evolutionary analysis of Australian *Pseudomys apodemoides* and *P. bolami* KoRV-related *gag pol* and *env* retroviral sequences.** Maximum likelihood phylogenies of regions of nucleotide sequence from the **A, D** *gag*, **B, E** *pol*, and **C, F** *env* genes overlapping contigs from **A-C** *P. apodemoides* and **D-F** *P. bolami*. All branches are scaled according to the number of nucleotide substitutions per site as indicated by the scale bars. Trees were rooted using the McERV (*Mus caroli* endogenous retrovirus) KC460271 or RfRV (*Rhinolophus ferrumequinum* retrovirus) JQ303225 sequences. Bootstrap support values are shown at the nodes.

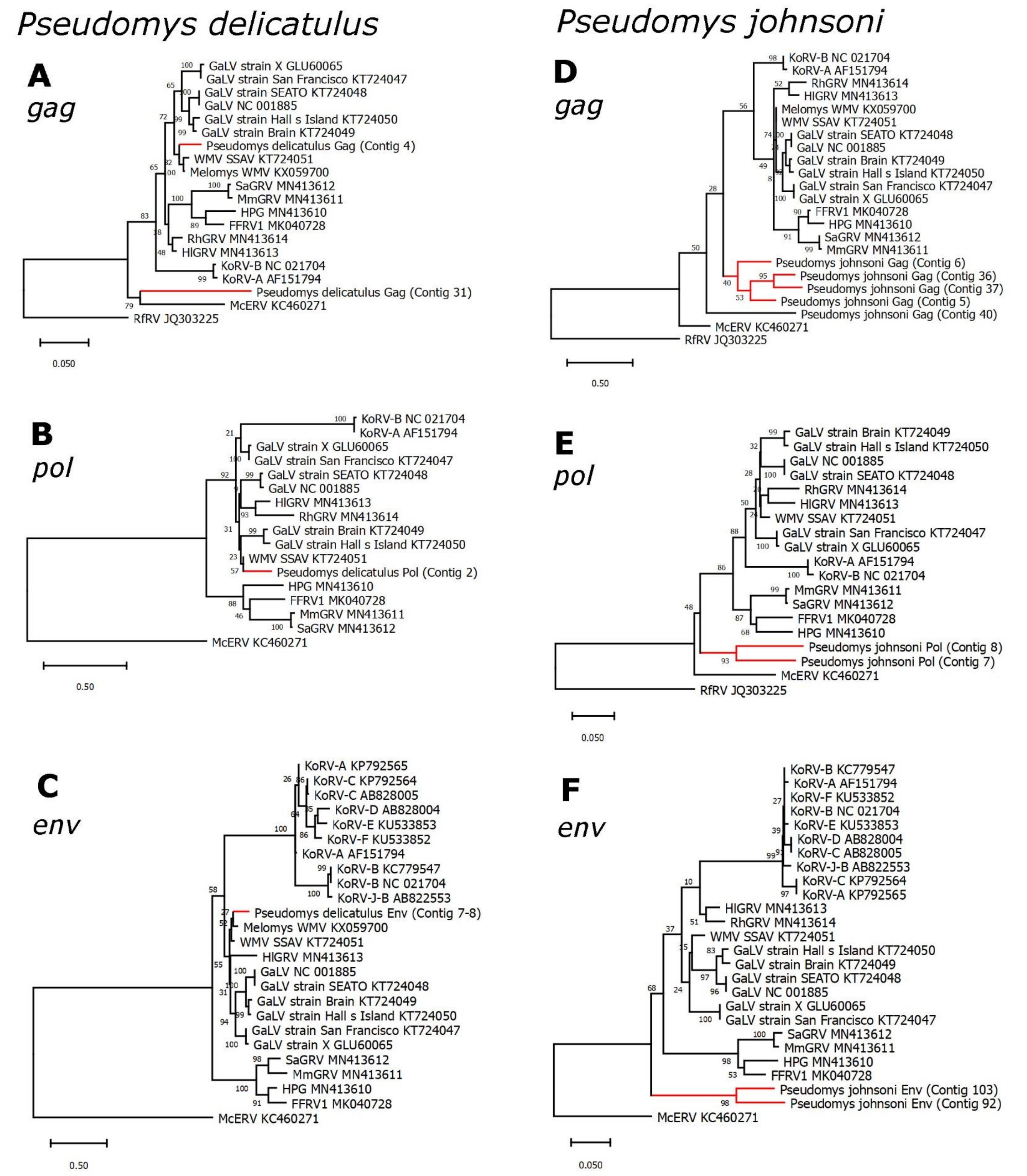


**SI Figure 3. Phylogenetic evolutionary analysis of Australian *Pseudomys delicatulus* and *P. johnsoni* KoRV-related *gag pol* and *env* retroviral sequences.** Maximum likelihood phylogenies of regions of nucleotide sequence from the **A, D** *gag*, **B, E** *pol*, and **C, F** *env* genes overlapping contigs from **A-C** *P. delicatulus* and **D-F** *P. johnsoni*. All branches are scaled according to the number of nucleotide substitutions per site as indicated by the scale bars. Trees were rooted using the McERV (*Mus caroli* endogenous retrovirus) KC460271 or RfRV (*Rhinolophus ferrumequinum* retrovirus) JQ303225 sequences. Bootstrap support values are shown at the nodes.

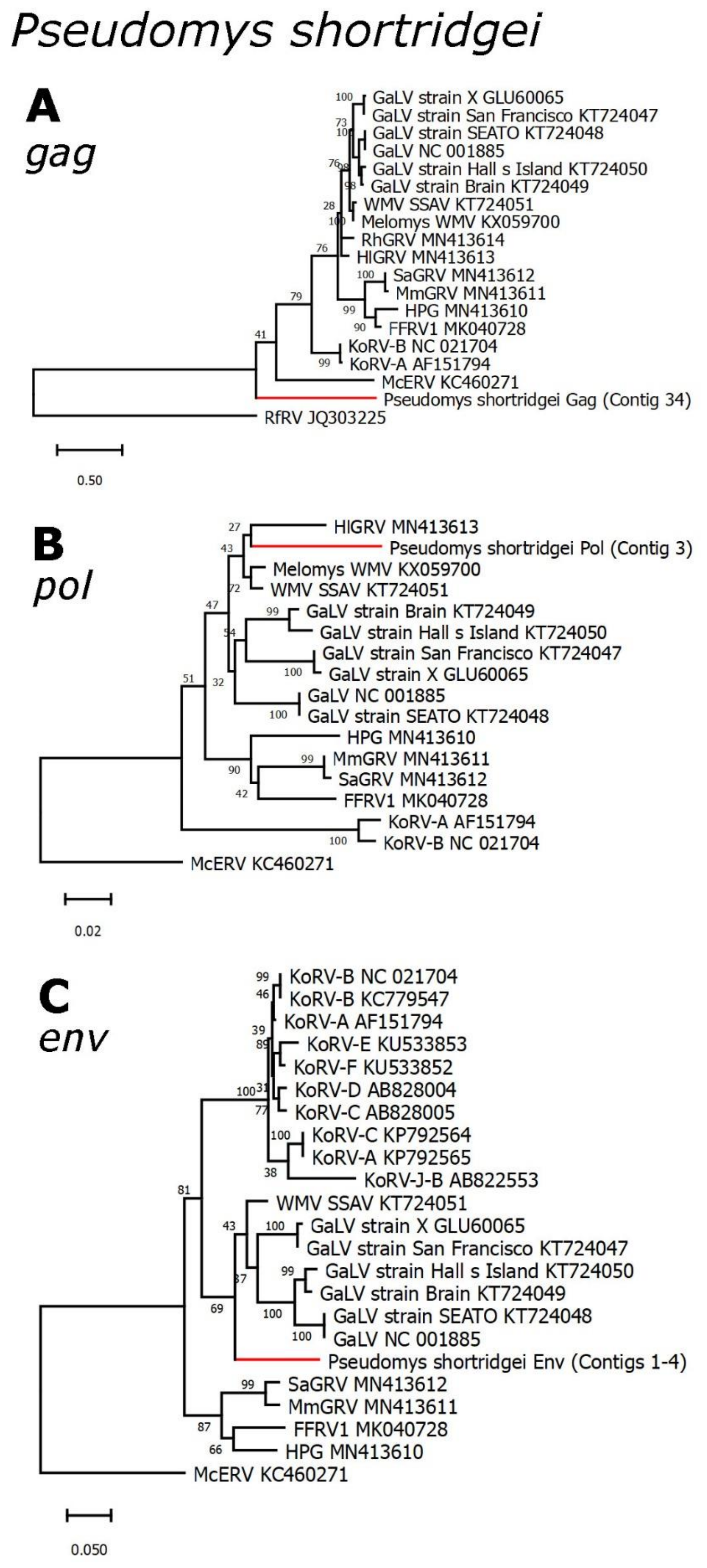


**SI Figure 4. Phylogenetic evolutionary analysis of Australian *Pseudomys shortridgei gag pol* and *env* KoRV-related retroviral sequences.** Maximum likelihood phylogenies of regions of nucleotide sequence from the **A** *gag*, **B** *pol*, and **C** *env* genes overlapping contigs from *P. shortridgei*. All branches are scaled according to the number of nucleotide substitutions per site as indicated by the scale bars. Trees were rooted using the McERV (*Mus caroli* endogenous retrovirus) KC460271 or RfRV (*Rhinolophus ferrumequinum* retrovirus) JQ303225 sequences. Bootstrap support values are shown at the nodes.

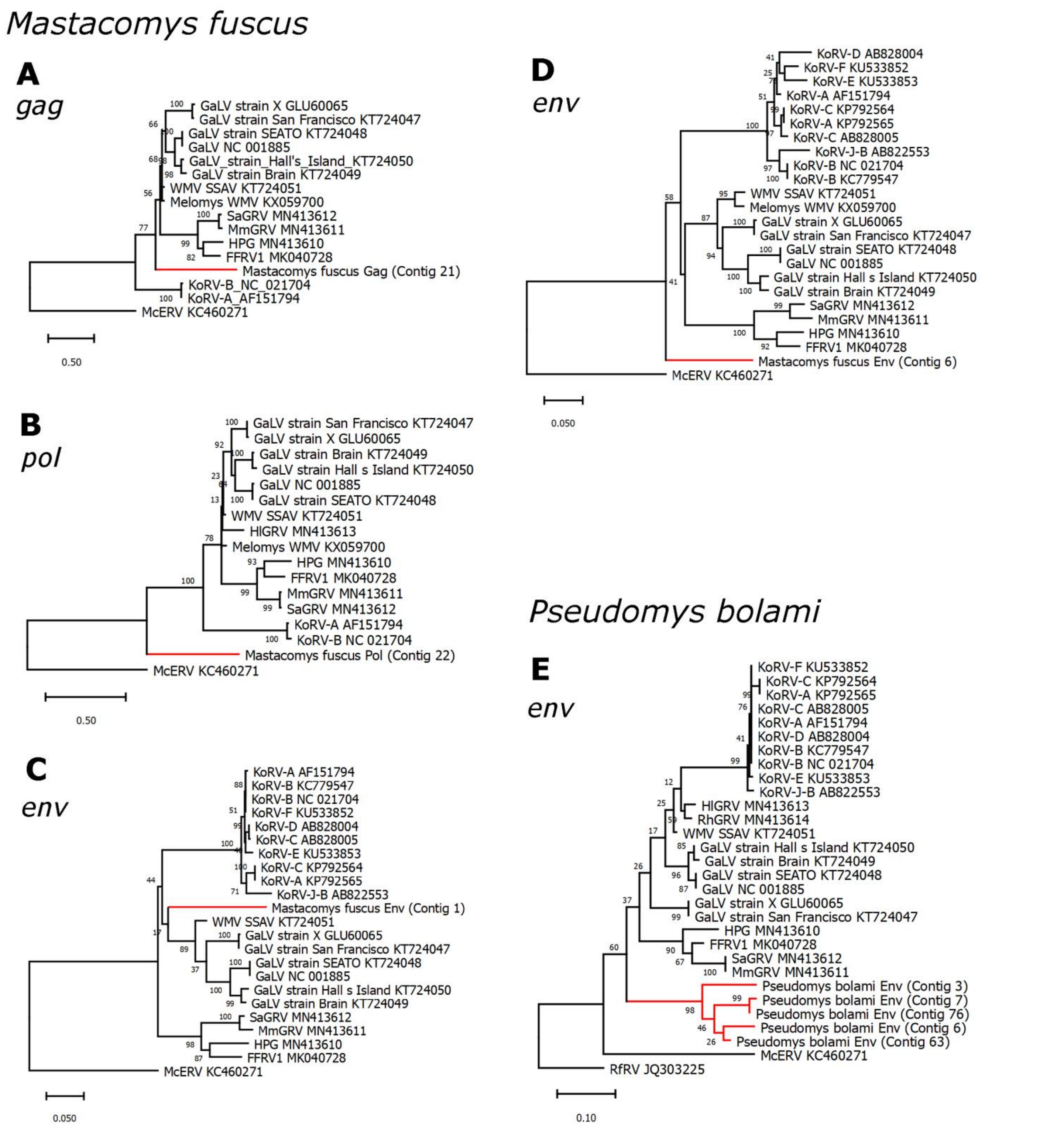


**SI Figure 5. Phylogenetic evolutionary analysis of additional *Mastacomys fuscus* and *Pseudomys bolami* KoRV-related retroviral sequences.** Maximum likelihood phylogenies of regions of nucleotide sequence from the **A** *gag*, **B** *pol*, and **C, D, E** *env* genes overlapping contigs from **A-D** *M. fuscus* and **E** *P. bolami*. All branches are scaled according to the number of nucleotide substitutions per site as indicated by the scale bars. Trees were rooted using the McERV (*Mus caroli* endogenous retrovirus) KC460271 or RfRV (*Rhinolophus ferrumequinum* retrovirus) JQ303225 sequences. Bootstrap support values are shown at the nodes.

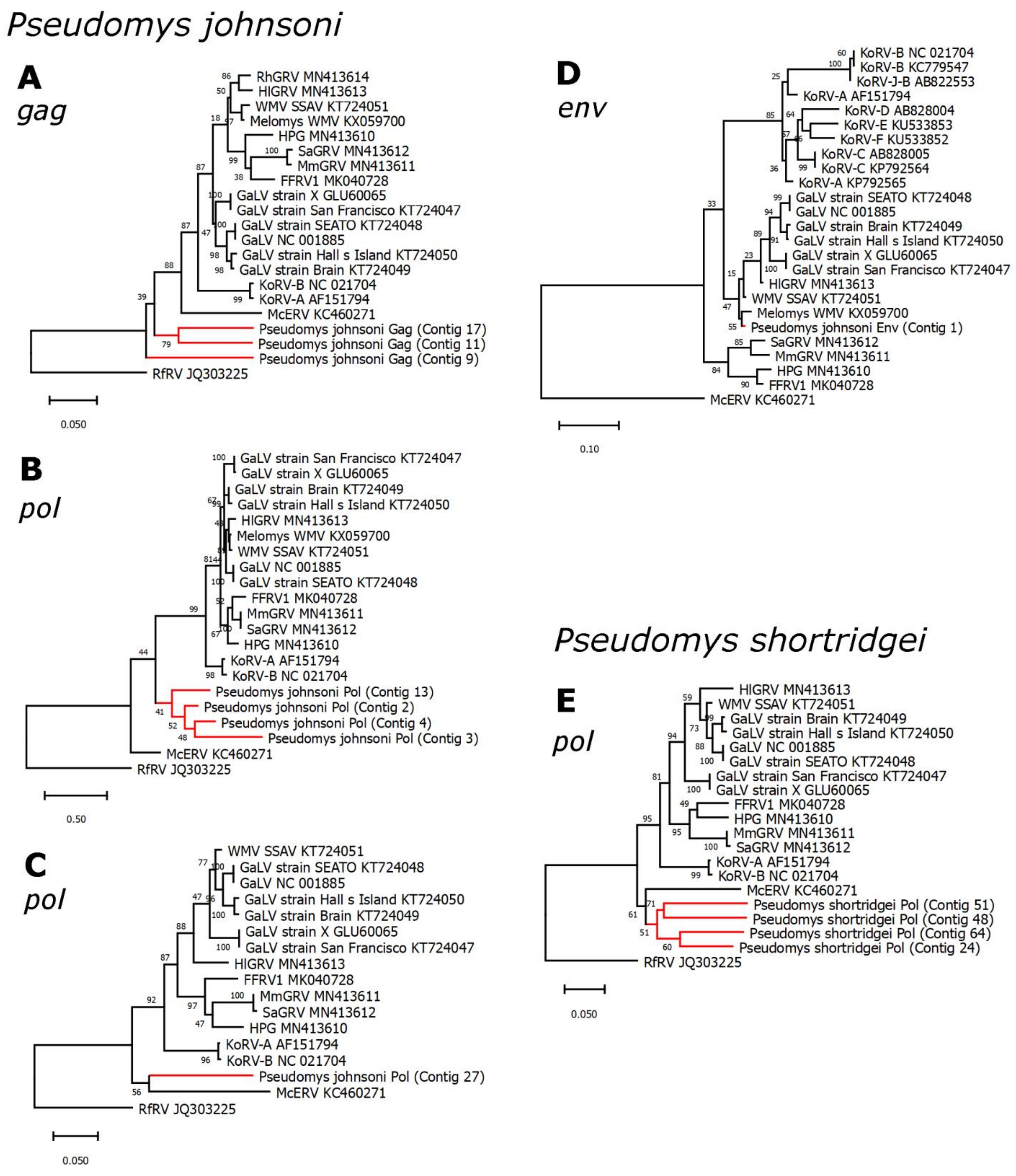


**SI Figure 6. Phylogenetic evolutionary analysis of additional *Pseudomys johnsoni* and *P. shortridgei* KoRV-related retroviral sequences.** Maximum likelihood phylogenies of regions of nucleotide sequence from the **A** *gag*, **B, C, E** *pol*, and **D** *env* genes overlapping contigs from **A-D** *P. johnsoni* and **E** *P. shortridgei*. All branches are scaled according to the number of nucleotide substitutions per site as indicated by the scale bars. Trees were rooted using the McERV (*Mus caroli* endogenous retrovirus) KC460271 or RfRV (*Rhinolophus ferrumequinum* retrovirus) JQ303225 sequences. Bootstrap support values are shown at the nodes.

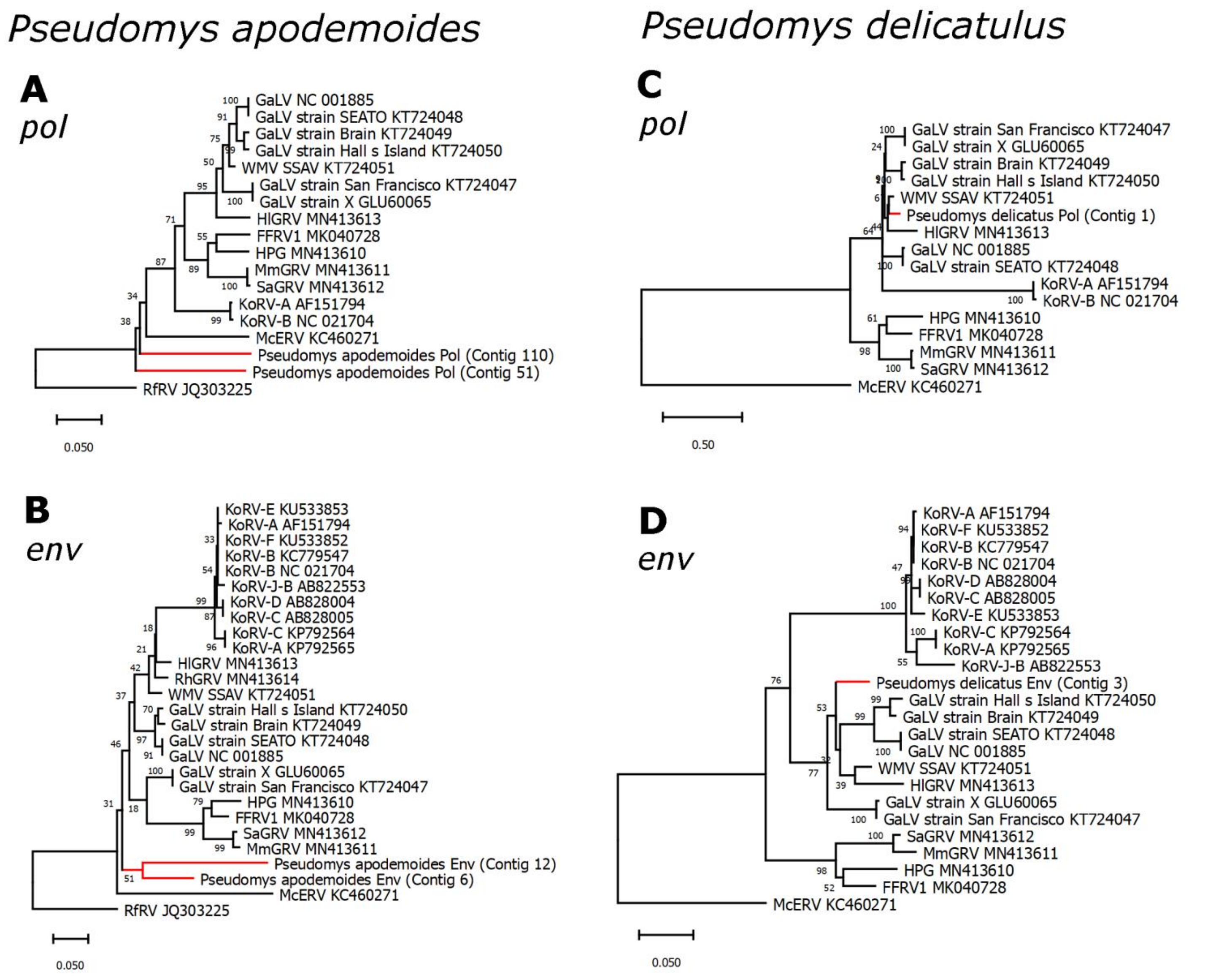


**SI Figure 7. Phylogenetic evolutionary analysis of additional *Pseudomys apodemoides* and *P. delicatulus* KoRV-related retroviral sequences.** Maximum likelihood phylogenies of regions of nucleotide sequence from the **A, C** *pol*, and **B, D** *env* genes overlapping contigs from **A, B** *P. apodemoides* and **C, D** *P. delicatulus*. All branches are scaled according to the number of nucleotide substitutions per site as indicated by the scale bars. Trees were rooted using the McERV (*Mus caroli* endogenous retrovirus) KC460271 or RfRV (*Rhinolophus ferrumequinum* retrovirus) JQ303225 sequences. Bootstrap support values are shown at the nodes.

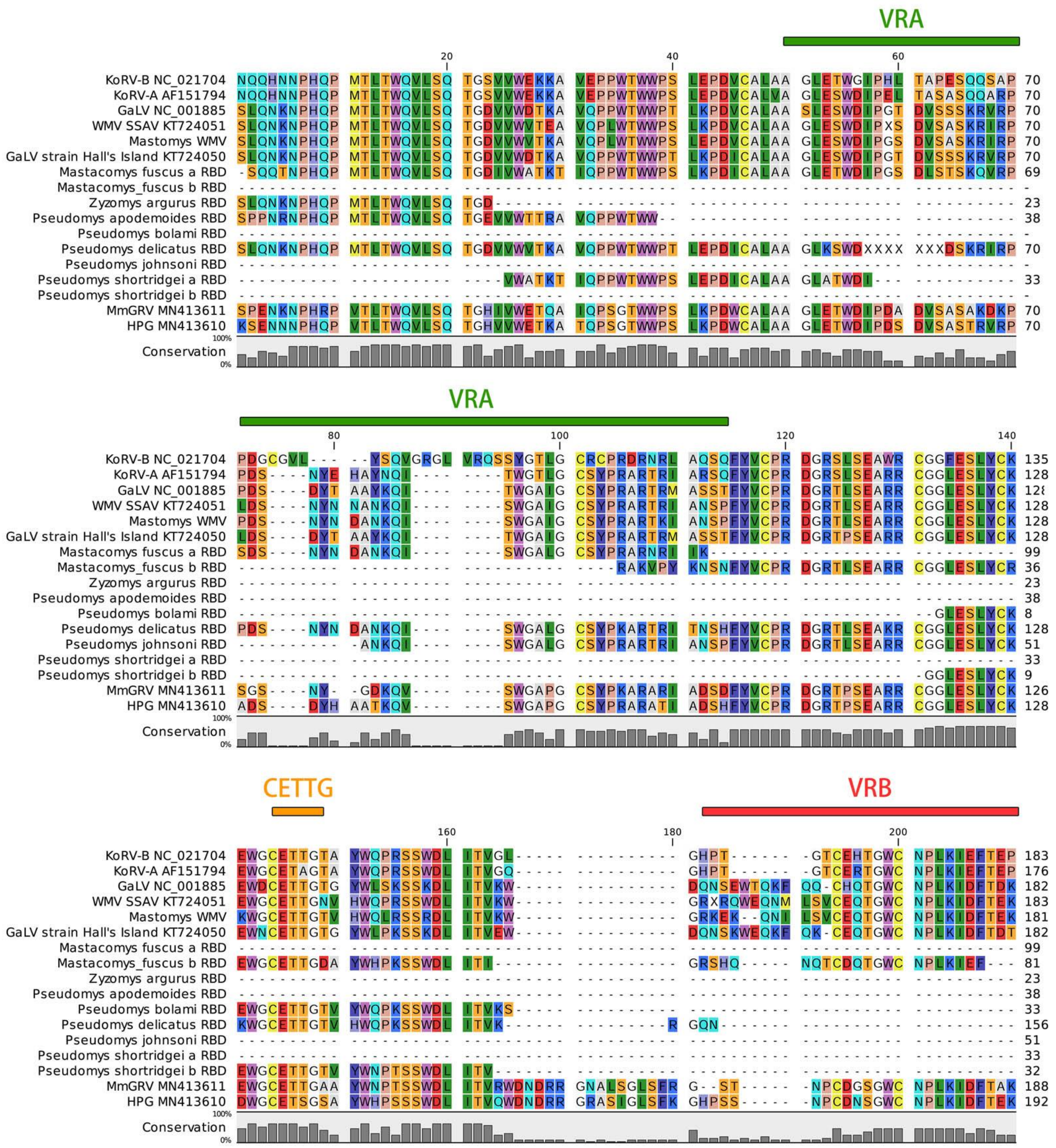


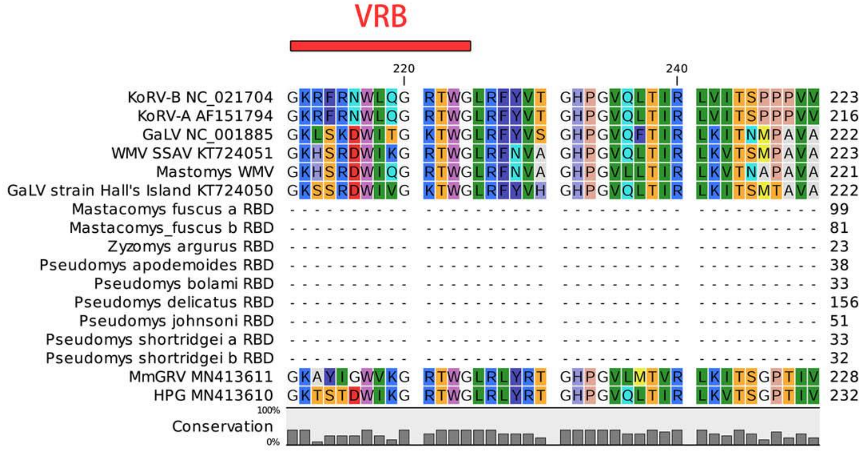


**SI Figure 8. Multiple sequence alignment of the receptor binding domain of novel rodent KoRV-related retroviruses.** The alignment was generated using MUSCLE (1). Novel sequences are compared against several known sequences. Highly variable regions A (VRA) and B (VRB) indicated by the green and red lines, respectively. The CETTG motif is denoted with an orange line. Due to the limited nature of the available sequences, not all sequences span the entire region.

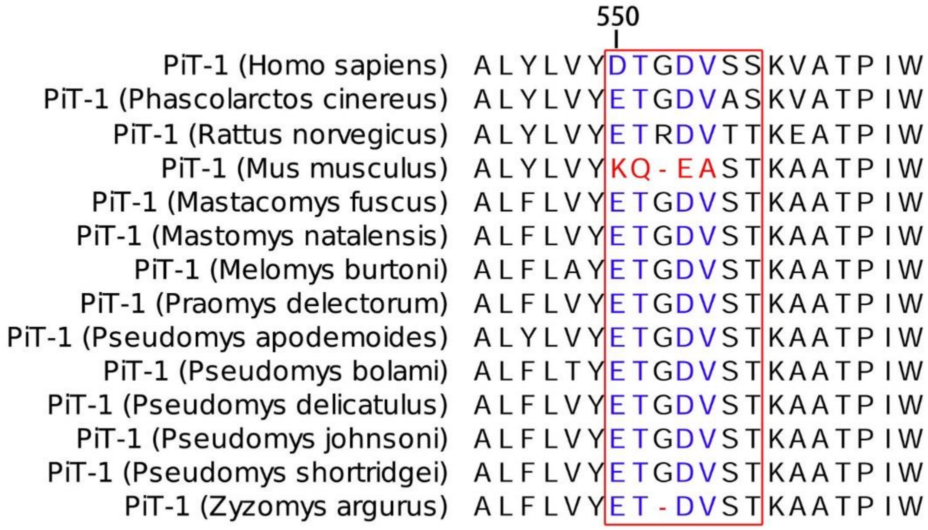


**SI Figure 9. Multiple sequence alignment of residues of the PiT-1 Region A of novel rodent hosts of KoRV-related retroviruses.** The Region A ‘permissivity’ motif of mammalian PiT-1 (SLC20A1) is shown in the red box (amino acid positions 550-557). Residues highlighted in blue and red denote residues in infection susceptible and resistant mammalian PiT-1 homologs, respectively (2).
